# Signal Versus Noise: Evaluating iNaturalist Photos as a Source of Quantitative Phenotypic Data in *Plethodon* Salamanders using Autoresearch and Agentic AI

**DOI:** 10.64898/2026.03.24.713936

**Authors:** Kyle A. O’Connell

## Abstract

Community-science platforms such as iNaturalist now contain tens of millions of georeferenced, photographically vouchered biodiversity records, yet extracting reliable quantitative measurements from opportunistic photographs remains methodologically challenging. Here, I evaluate the signal-to-noise ratio of iNaturalist photos for phenotyping *Plethodon* salamanders across two trait classes: continuous dorsal brightness (a proxy for ecogeographic clines predicted by Gloger’s rule and the thermal melanism hypothesis) and discrete color morph frequency in *P. cinereus*. I optimized a color-extraction pipeline using an agent-guided parameter search adapted from the *autoresearch* framework (Karpathy 2026; Schmidgall et al. 2025), exploring crop fraction, color space, normalization, and quality-control thresholds across 50 bounded micro-experiments. Applying the production HSV pipeline to 103,653 observations of 34 species, I found negligible geographic structure in dorsal brightness (R^2^ = 0.001), even within *P. cinereus* alone (n = 71,627). Variance decomposition showed that photographer identity explains 23.3% of brightness variance, geography 5.1%, species 1.6%, and time of day 0.3%, with 69.7% residual. In contrast, a hue-threshold morph classifier recovered a significant geographic signal in red-back frequency (R^2^ = 0.008, p < 0.001), 7× stronger than the brightness result, though still weaker than the supervised CNN of Hantak et al. (2022; pseudo-R^2^ ≈ 0.04). These results indicate that citizen-science photographs are poorly suited to continuous quantitative phenotyping under current collection conditions, whereas discrete categorical traits remain recoverable with appropriate classifiers. The autoresearch loop clarified the failure mode: no tested parameter configuration recovered a meaningful brightness signal from a dataset dominated by observer effects.

## Introduction

Color is among the most tractable phenotypic traits in ecology and evolutionary biology. It is directly observable, ecologically meaningful, and predictably distributed across geography in ways that link physiology, behavior, climate, and natural selection (Endler 1980; McLean and Stuart-Fox 2014). Two classic ecogeographic rules make explicit, opposing predictions about how coloration should vary with latitude and climate in terrestrial ectotherms. Gloger’s rule predicts darker pigmentation in warmer, more humid environments, consistent with adaptive responses to UV radiation, desiccation, and parasitism (Delhey 2019). The thermal melanism hypothesis predicts darker coloration at higher latitudes and elevations, where darker bodies absorb solar radiation more efficiently and support faster warming in cool environments (Clusella-Trullas et al. 2007). In a genus of animals spanning ∼25° of latitude across eastern North America, the direction of any observed latitudinal brightness gradient would immediately inform which selective pressure dominates.

*Plethodon* (family Plethodontidae) is the most species-rich vertebrate genus in eastern North America, comprising ∼55 species of lungless salamanders that occupy forest floor microhabitats from Florida to Newfoundland and from sea level to the high Appalachians. The genus is exceptionally well studied from community ecology and biogeographic perspectives (Hairston 1951; Jaeger 1971; Highton 1995), but quantitative range-wide analysis of dorsal coloration has been limited by the difficulty of obtaining standardized photographs across the full geographic range. iNaturalist (Di Cecco et al. 2021; iNaturalist contributors 2026) now hosts >154,000 Research Grade *Plethodon* observations, each linked to a georeferenced photograph, representing orders of magnitude more geographic coverage than any museum collection or directed field survey.

The central challenge in using this resource for quantitative phenotyping is that iNaturalist photographs are taken by thousands of observers under heterogeneous conditions, with uncontrolled lighting, camera settings, and subject positioning. Extracting measurements that reflect biological rather than photographic variation therefore requires a pipeline that is robust to image heterogeneity and explicitly evaluated for its ability to recover known signals. This evaluation step, namely determining *what* the photos can and cannot reveal, is not yet routine in citizen-science phenotyping studies.

Recent work has begun to address this gap for discrete phenotypic classes. Hantak et al. (2022) demonstrated that an ensemble convolutional neural network (CNN) trained on 4,000 human-labeled iNaturalist images could classify the striped/unstriped color polymorphism in *P. cinereus* with ∼98% accuracy across 20,318 images, and used the resulting annotations to document climatic niche associations between morphs at range-wide scale (pseudo-R^2^ ≈ 0.04). The success of this approach depended on supervised training with expert-labeled data and a whole-image deep learning architecture capable of implicitly localizing the relevant dorsal region. Whether simpler, unsupervised photometric methods — mean pixel brightness, hue, or saturation extracted from a central crop — can recover comparable signals remains unknown.

For continuous traits, the question is more fundamental: are the photos informative at all? Observer-level heterogeneity in exposure, flash use, and photo composition creates additive noise that may overwhelm subtle quantitative gradients. McCormick and Riley (2025) documented systematic observer bias in the same system: in a concurrent field survey in New Brunswick, Canada, 95.9% of encountered striped-morph individuals were striped, whereas only 83.2% of iNaturalist observations were striped — a significant overrepresentation of rarer morphs in the community science dataset (χ^2^ = 11.92, p < 0.01). At the level of specific rare morphs, the unstriped form was observed at 6.6% on iNaturalist versus 3.3% in field surveys (χ^2^ = 5.83, p = 0.02). This suggests that observer selection preferences distort even categorical measurements. Whether analogous biases distort continuous brightness measurements has not been formally evaluated.

Here, I address these questions using three complementary analyses. First, I optimize and benchmark a color extraction pipeline for iNaturalist *Plethodon* photographs using a bounded, agent-guided parameter search adapted from Karpathy’s (2026) *autoresearch* framework and related LLM-agent research-assistant workflows (Schmidgall et al. 2025), which treat parameter optimization as a logged sequence of micro-experiments evaluated against a composite geographic signal score. Second, I apply a stable production extraction pipeline to 103,653 observations and use the pilot-optimized alternative as a sensitivity check on the validation subset. Third, I decompose brightness variance into biological and photographic sources using hierarchical ICC analysis and a linear mixed model, and compare the resulting signal-to-noise profile against a simple hue-threshold morph classifier applied to the same photographs. My goal is to provide an explicit empirical characterization of what iNaturalist photographs can and cannot reveal about phenotypic variation in a well-studied salamander clade — and to demonstrate a reproducible, computationally formalized approach to optimizing extraction pipelines for citizen science image data.

## Methods

### Study System and Data Source

All observations were obtained from iNaturalist through the iNaturalist API (Di Cecco et al. 2021; iNaturalist contributors 2026). I downloaded metadata for 103,669 Research Grade observations spanning 34 species and covering the eastern and Pacific *Plethodon* radiations across latitudes 25–50°N. Research Grade status requires that an observation be accompanied by a usable photograph and that species identity be confirmed by a minimum of two independent community reviewers. Preprocessing included removal of observations with missing or obscured geographic coordinates, deduplication of near-identical records within 100 m, standardization of observation fields, and flagging of potential out-of-range occurrences based on known species distributions. After preprocessing and image-level quality-control filtering (brightness and entropy thresholds; see Color Feature Extraction below), the final analysis dataset comprised 103,653 observations — a net loss of only 16 records, reflecting the permissive nature of the automated QC thresholds documented in the crop quality audit. Cleaned observations were assigned to H3 hexagonal grid cells at resolution 5 (∼252 km^2^ per cell) using the h3-py library (Brodsky 2018). This spatial binning unit provides a geographically consistent footprint for estimating within-location variance and supports the geographic signal score used in pipeline optimization.

From the cleaned observation table, I generated a photo manifest containing the first available image URL for each record. Images were downloaded at a rate-limited throughput of ∼10 requests/second using a shared token-bucket limiter across six concurrent threads, with per-request retry logic and checkpoint recovery. Downloaded images were stored by observation identifier to maintain a one-to-one mapping between extracted color measurements and source records.

### Agent-Guided Optimization of the Color Extraction Pipeline

The image analysis workflow was framed as a constrained optimization problem adapted from the *autoresearch* framework (Karpathy 2026), in which parameter search is structured as a logged sequence of bounded micro-experiments rather than ad hoc manual tuning. An LLM agent (claude-opus-4-6) iteratively proposed one parameter change at a time to a restricted configuration dictionary, executed the revised pipeline on a fixed validation subset of ∼859 photographs, evaluated the result against a predefined composite score, and retained or reverted the change before proposing the next experiment.

The agent was constrained to modify only the following parameters:

- `central_crop_fraction` [0.15–0.7]: Fraction of image area retained in the centered dorsal crop
- `color_space` [HSV, LAB, RGB]: Color representation used for brightness extraction
- `normalize_brightness` [bool]: Whether to apply per-channel histogram equalization before extraction
- `background_mask` [none, green_threshold, saturation_threshold]: Optional pixel exclusion strategy
- `min_brightness`, `max_brightness`, `min_entropy`: Quality-control rejection thresholds
- `percentile_trim` [0–10%]: Fraction of extreme pixel values excluded from mean calculation

The composite optimization score rewarded geographic signal while penalizing local noise:

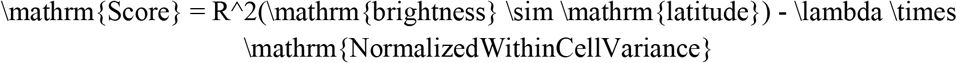

where $\lambda = 0.5$. Each experiment returned extracted brightness values linked to observation coordinates and H3 cells; from these, I computed the latitude regression R^2^ and mean within-cell brightness variance across cells with at least three observations. All experiments were logged with their full parameter configuration, score, regression statistics, and QC summary.

The pilot optimization loop ran for 50 iterations on 859 photographs from an initial test dataset. I then deployed the same workflow on a Google Cloud Platform VM (n2-highmem-32, 32 vCPUs, 256 GB RAM, 200 GB SSD boot disk, Debian 12, `us-east1-b`) to handle large-scale photo download, additional optimization, and production extraction. The VM operated autonomously through a startup script with no interactive access: it cloned the repository, installed dependencies, downloaded the full photo corpus, ran the optimization loop, performed final extraction, uploaded results, and self-terminated. An initial spot-instance deployment was preempted after 2.5 hours during photo download; the final production run therefore used standard on-demand provisioning (∼$0.33/hr spot vs. ∼$2.10/hr on-demand for n2-highmem-32), with an estimated total cost of ∼$21 for ∼10 compute-hours. All local preprocessing, downstream statistical analysis, and figure generation were performed on a MacBook Air (M3, 16 GB RAM, macOS 26.3).

### Color Feature Extraction

The pilot autoresearch loop identified CIE L\*a\*b\* color space and histogram normalization as the largest improvements to the composite score. For the primary manuscript analyses, however, I report the stable HSV production extraction (40% central crop, no normalization) that had already completed across the full dataset when results were frozen for analysis. The pilot-optimized LAB configuration is therefore treated as a validation-subset sensitivity check rather than as the production pipeline. Although the optimized configuration substantially reduced within-cell variance on the validation subset, both HSV and LAB pipelines led to the same qualitative conclusion: geographic brightness signal remained negligible relative to photographic noise. All reported brightness values therefore use HSV V-channel measurements on a 0–255 scale.

For each downloaded photograph, I extracted dorsal color features from a centered crop using the following procedure: (1) load image as RGB using Pillow; (2) extract the central 40% × 40% fraction; (3) convert to HSV using scikit-image (H in degrees 0–360, S and V on 0–255 scales); (4) compute Shannon entropy of the full crop as an image-quality indicator; (5) apply quality-control filters (brightness □ [15, 245], entropy ≥ 4.0) and exclude images that fail. Retained images contributed five metrics per observation: `mean_brightness` (V channel), `mean_hue` (H channel), `mean_saturation` (S channel), `entropy`, and image dimensions. After QC filtering and merging to observation metadata (lat, lon, species, H3 cell, observer, date), the final analysis dataset comprised 103,653 observations.

### Geographic Brightness Analysis

I tested for geographic clines in dorsal brightness using ordinary least squares (OLS) regression of `mean_brightness` against latitude and longitude (brightness ∼ lat + lon), implemented in statsmodels. I fit this model (1) across all 34 species simultaneously (n = 103,653) and (2) for *P. cinereus* alone (n = 71,627), the most-photographed species and the one for which observer bias and morph composition are best characterized. At the cell level, I aggregated mean brightness per H3 res-5 cell and re-fit the model on cell means (n = 6,474 cells). A positive latitude coefficient would indicate brighter dorsal coloration at higher latitudes (consistent with Gloger’s rule); a negative coefficient would indicate darker coloration (consistent with thermal melanism). I report R^2^, adjusted R^2^, F-statistic p-value, and regression coefficients with standard errors. Cross-species heterogeneity in brightness was tested with a Kruskal– Wallis test across all species with ≥10 observations.

### Variance Decomposition

To partition brightness variation into biological versus photographic sources, I computed intraclass correlation coefficients (ICC) for four grouping variables: species identity (34 groups), individual observer (`user_id`, 33,496 groups), hour of day (24 groups, derived from the timezone-aware `observed_on` timestamp), and H3 res-5 geographic cell (6,474 groups). The ICC(1) estimator for each variable was:

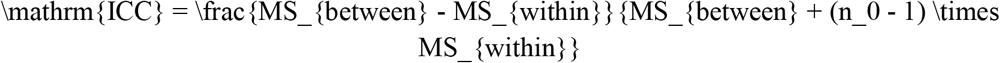

where $n_0$ is the mean group size and $MS_{between}$/$MS_{within}$ are the between- and within-group mean squares from a one-way ANOVA. I also fit a linear mixed model (statsmodels MixedLM, REML) with `user_id` as a random intercept and latitude, longitude, species code, hour of day, image entropy, and aspect ratio as fixed effects, to estimate the proportion of total variance attributable to the photographer random effect. Feature engineering added `hour_of_day` (extracted from `observed_on`), `aspect_ratio` (width/height), and `log_obs_per_user` (log of observations per observer, as an experience proxy) to the analysis dataset. ICC values were floored at zero; residual variance was estimated as 100% minus the sum of all ICC terms.

### Color Morph Classification and Geographic Validation

As a positive control for geographic signal detection, I applied a hue-threshold classifier to the 71,627 *P. cinereus* observations to distinguish red-back (striped) from lead-back (unstriped) morphs. Following the documented pigment biology of this polymorphism (Lotter and Scott 1977), I classified observations as red-back if `mean_hue` fell within the warm range (0–60° or 300–360°) and `mean_saturation` ≥ 30 (on a 0–255 scale). Observations with `mean_saturation` < 30 were classified as lead-back regardless of hue; observations with hue in the clearly cool range (90–270°) and saturation ≥ 30 were classified as lead-back; all remaining observations were classified as ambiguous and excluded from geographic analysis. These conservative thresholds prioritize classification confidence over coverage, retaining only observations with unambiguous photometric evidence of morph identity.

Classified observations (excluding ambiguous) were aggregated per H3 res-5 cell to compute red-back morph frequency (n_redback / n_total). I fit an OLS model of red-back frequency ∼ mean cell latitude + mean cell longitude using cells with ≥5 classified observations (n = 1,776 cells). The resulting R^2^ was compared directly to the brightness R^2^ as the primary signal-versus-noise contrast. All analyses were implemented in Python using pandas, numpy, scipy, and statsmodels; figures were generated with matplotlib and seaborn. Code and derived analysis outputs are available at https://doi.org/10.5281/zenodo.19050224.

### Crop Quality Validation

To assess whether the center-crop protocol reliably isolates dorsal salamander body pixels, I conducted a manual audit of 200 randomly sampled locally cached photographs (seed 20260323). Each crop was scored by a single rater using a four-category rubric: (1) *good* — crop contains mostly salamander trunk/dorsal pixels, acceptable for brightness extraction; (2) *partial* — salamander present but substantial non-body content remains; (3) *fail* — crop is not usable dorsal body pixels; (4) *in_hand* — hand or glove is a major crop component. I additionally recorded `brightness_usable` (yes/no/unclear) and `morph_usable` (yes/no/unclear) per image, derived conservatively from crop-quality category. This audit was designed as a single-rater protocol failure-rate estimate, not an inter-rater validation study.

Audit results were used to compute: (a) concordance between the automated `passed_qc` flag and manual crop-quality labels; (b) a test of whether poor-quality crops are geographically structured (Mann– Whitney U comparing latitude distributions of `good` versus `not-good` observations); and (c) two sensitivity regression analyses. Path A restricted to `brightness_usable = yes` observations from the audit-matched parquet subset (n = 75). Path C trained a logistic regression classifier (L2 regularization, C = 1.0) on five features available in the merged parquet (`mean_brightness`, `mean_hue`, `mean_saturation`, `entropy`, `aspect_ratio = width/height`) using the 195 audit-labeled observations, evaluated by 5-fold cross-validation, and applied to the full 103,653-observation dataset. Sensitivity regressions on the Path C–filtered subset were compared to full-dataset results, and the species composition of predicted-good versus predicted-bad subsets was inspected to assess whether the classifier introduced morph-sampling bias.

## Results

### Agent-Guided Pipeline Optimization

The pilot autoresearch loop completed 50 experiments on 859 photographs, of which 12 (24%) were accepted as improvements to the composite score. The baseline configuration (40% central crop, HSV color space, no normalization, no masking) achieved a composite score of −0.257, driven primarily by high within-cell brightness variance (mean = 1,101). Over the course of accepted experiments, the composite score improved to +0.010 — a +0.267 absolute improvement — while the latitude R^2^ remained stable at 0.018 and within-cell variance fell 97% to 33 (Table 1).

**Table 1.** Pilot autoresearch loop summary (composite score uses λ = 0.5).

| Parameter | Baseline | Optimized |
| --- | --- | --- |
| Color space | HSV | CIE L*a*b* |
| Crop fraction | 0.40 | 0.64 |
| Histogram normalization | No | Yes |
| Background masking | None | None |
| Percentile trim | 0% | 4.7% |
| Composite score | -0.257 | +0.010 |
| $R^2$ (brightness ~ latitude) | 0.018 | 0.018 |
| Within-cell variance | 1,101 | 33 |

The most consequential single change was histogram normalization (+0.163 to composite score), which standardized variable exposure across photos. Switching from HSV to CIE L\*a\*b\* color space (+0.058) improved the perceptual uniformity of brightness estimates, and a larger crop fraction (+0.021) captured more dorsal surface area. Background-masking strategies (green-channel threshold, saturation threshold) were explored but never improved the score and were ultimately excluded from the final configuration. The 24% acceptance rate is consistent with a well-constrained search: most proposed changes either had negligible effect or degraded the within-cell noise metric.

### Geographic Brightness Analysis

Applying the HSV production extraction pipeline to 103,653 observations across 34 species, I found negligible geographic structure in dorsal brightness. The OLS model brightness ∼ latitude + longitude explained R^2^ = 0.001 across all species (adj. R^2^ = 0.001, F-test p < 0.001; Table 2). The result held within *P. cinereus* alone (n = 71,627; R^2^ = 0.001, p < 0.001), where species composition is held constant. At the H3 cell level (n = 6,474 cells), R^2^ = 0.003 across all species and R^2^ = 0.001 for *P. cinereus* cells. Estimated regression coefficients were near zero (latitude: β ≈ 0.04 V/degree, longitude: β ≈ 0.01 V/degree) with confidence intervals spanning both positive and negative values. A Kruskal–Wallis test confirmed that species differ significantly in mean brightness (H = 830.1, p = 1.56 × 10□^1^□□), indicating that substantial cross-species signal exists in the dataset — it is the *within-species, across-geography* gradient that is absent. The regression panels, range-wide brightness map, and species-comparison plot (Figures 2-4) all reinforce the same pattern: substantial between-species spread but no clear within-species geographic cline.

**Table 2.** OLS brightness ∼ latitude + longitude results.

| Dataset | n | $R^2$ | adj. $R^2$ | F p-value |
| --- | --- | --- | --- | --- |
| All species (observation level) | 103,653 | 0.0011 | 0.0011 | < 0.001 |
| *P. cinereus* only (observation level) | 71,627 | 0.0011 | 0.0011 | < 0.001 |
| All species (cell level, $n \geq 3$) | 6,474 | 0.003 | 0.003 | < 0.001 |

**Figure 1.**
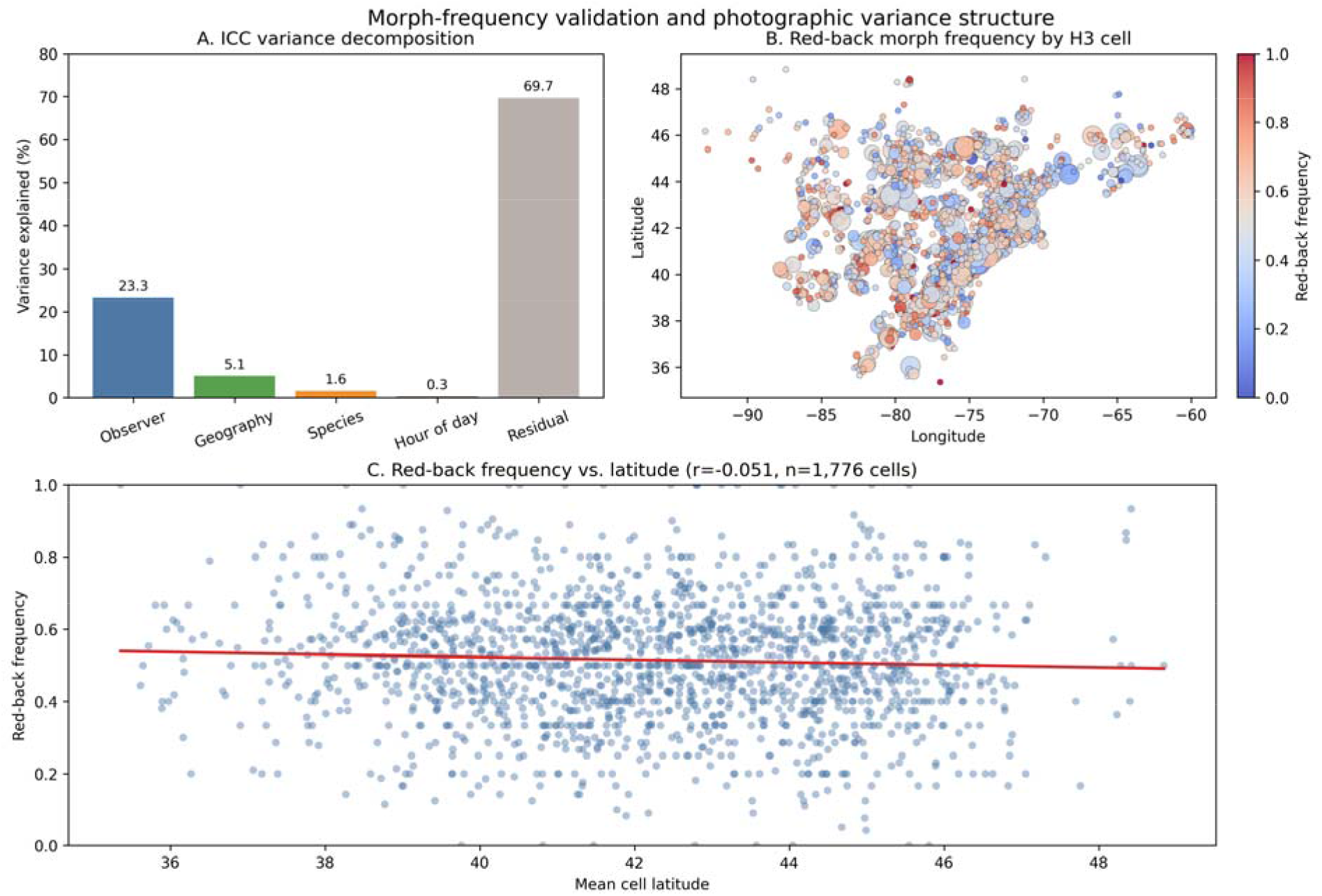
Composite summary of variance decomposition and morph-frequency validation. (A) ICC variance decomposition across observer, geography, species, hour-of-day, and residual sources. (B) H3-cell map of classified red-back morph frequency in *Plethodon cinereus*. (C) Red-back morph frequency as a function of mean cell latitude for cells with at least five classified observations.

**Figure 2.**
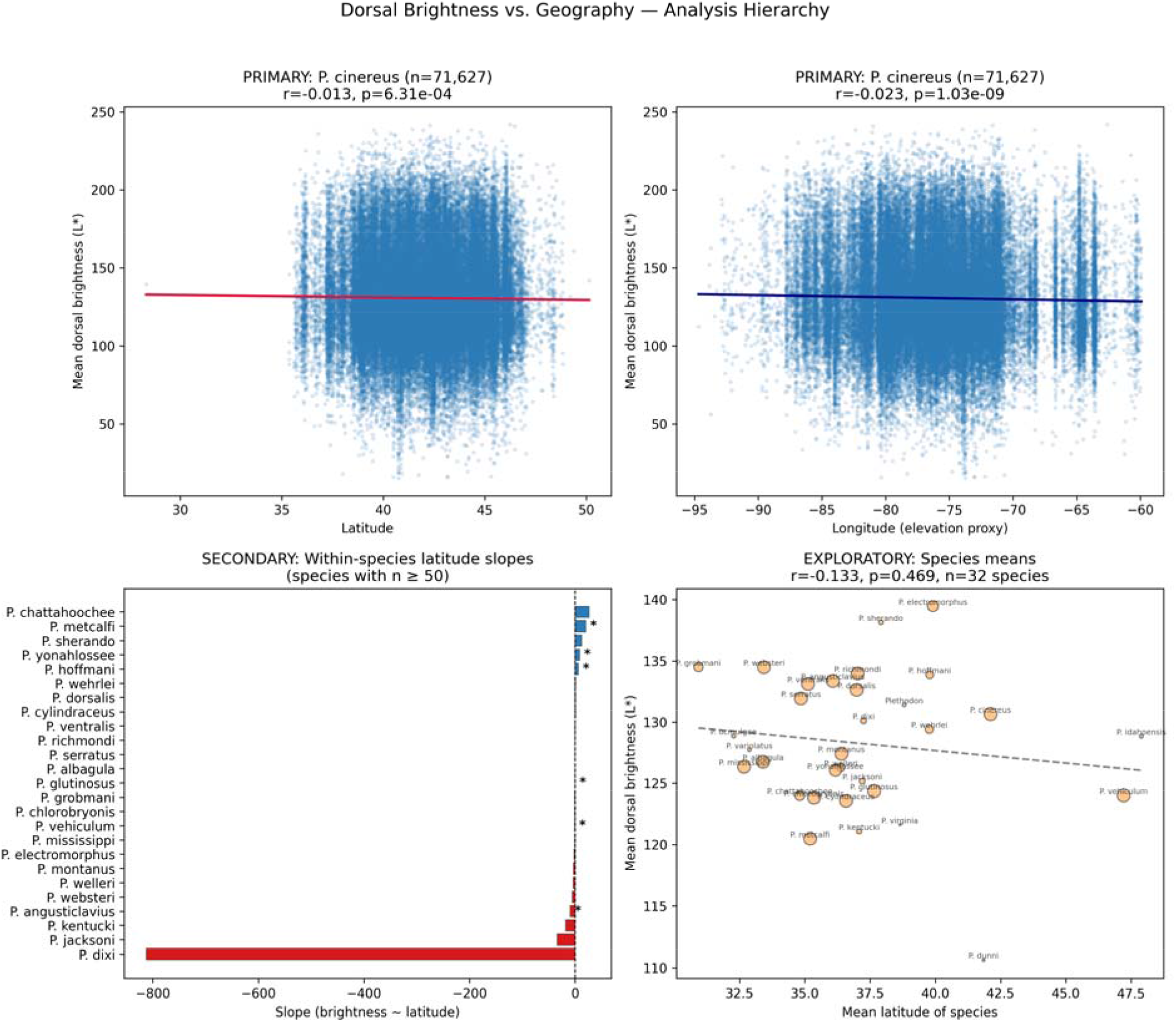
Dorsal brightness versus geography across analytic levels, including observation-level regressions, within-species slopes, and species means.

**Figure 3.**
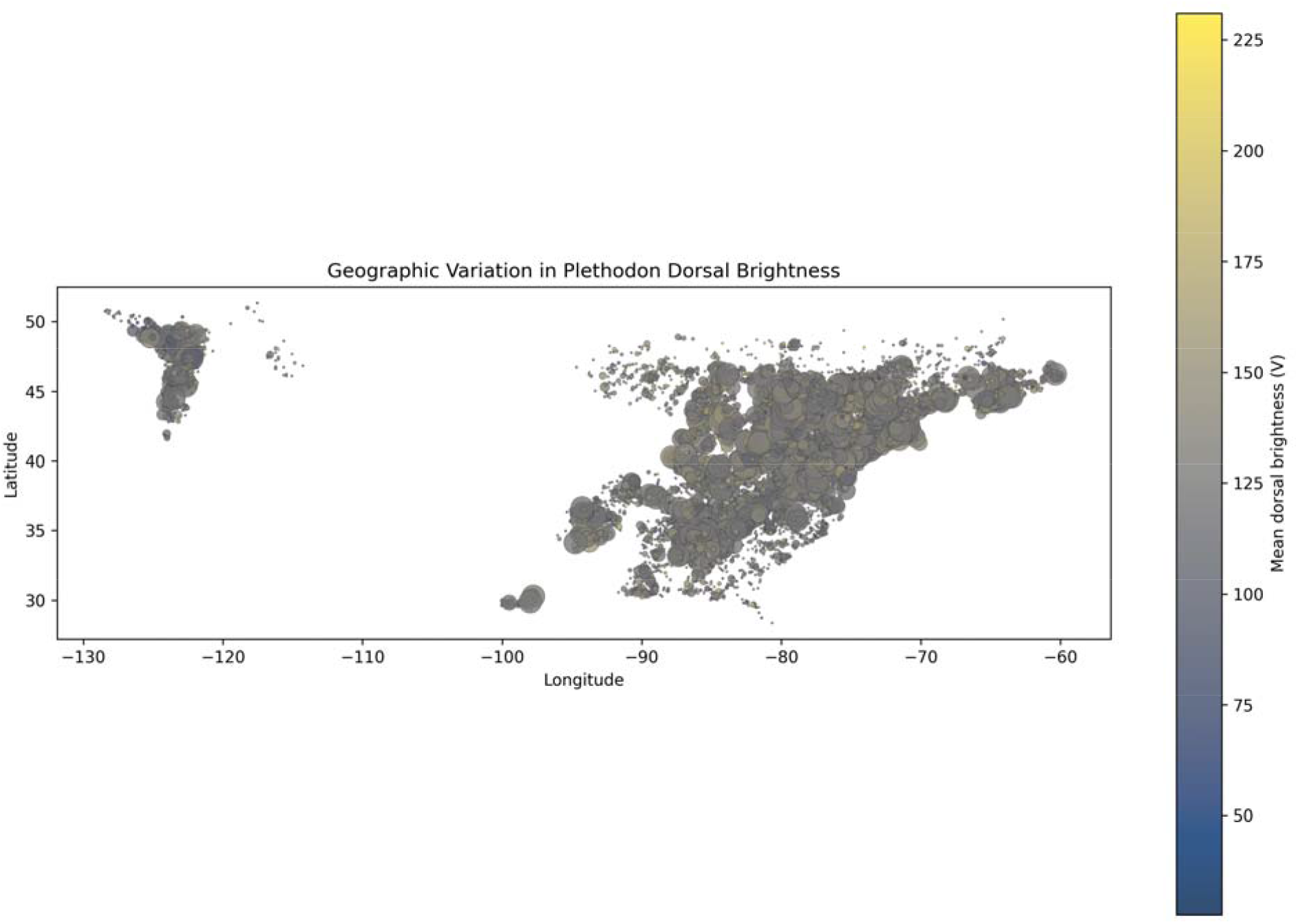
Geographic variation in *Plethodon* dorsal brightness across the study extent.

**Figure 4.**
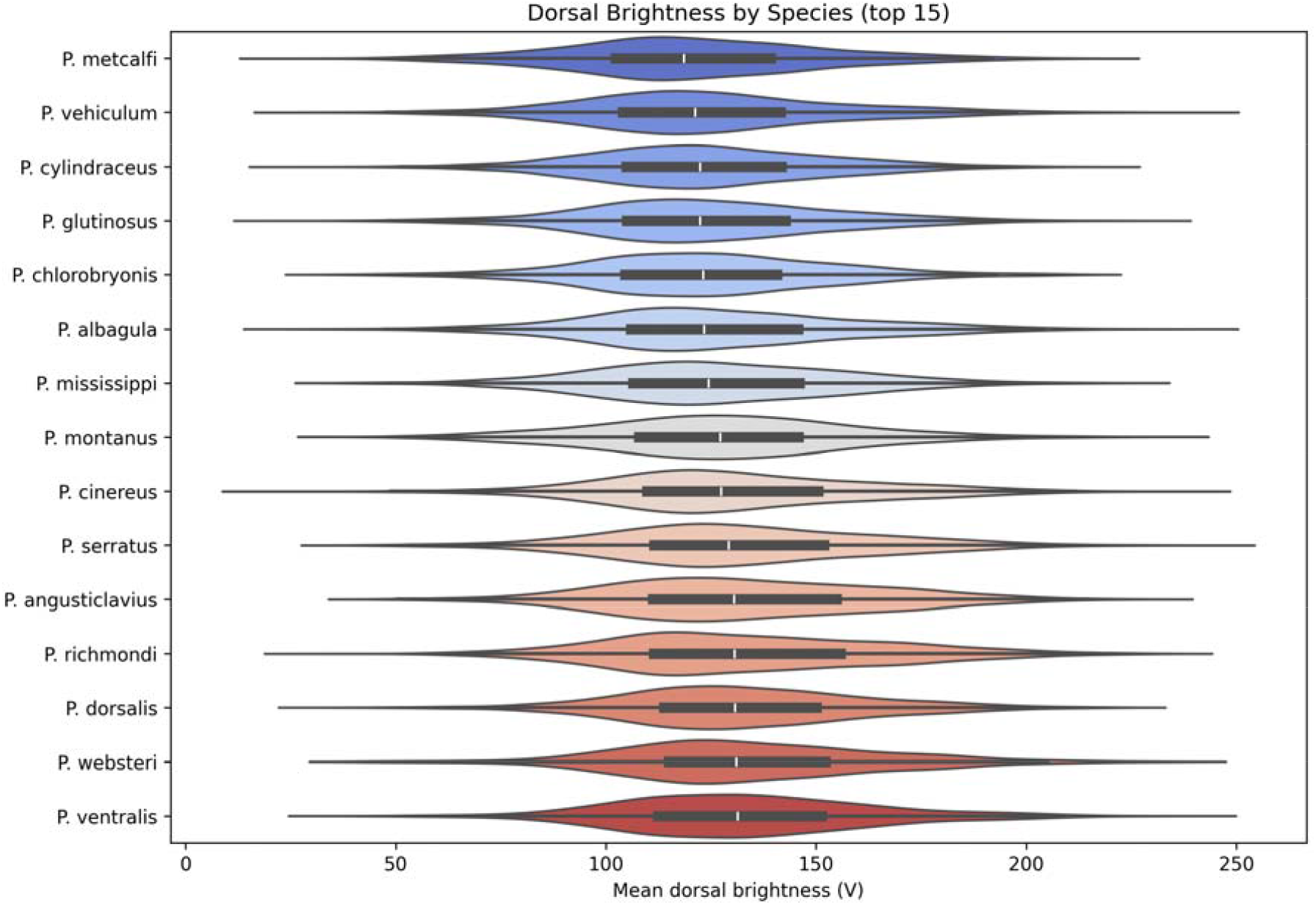
Distribution of dorsal brightness values for the 15 most-observed *Plethodon* species.

### Variance Decomposition

ICC analysis partitioned brightness variance into four identifiable sources (Figure 1A, Table 3). Observer identity (user_id) showed the largest ICC (0.233), indicating that individual photographers account for 23.3% of total brightness variance — more than all other modeled sources combined. Geographic cell (ICC = 0.051) explained 5.1%, reflecting a mixture of real spatial structure and spatially correlated photography conditions. Species identity explained 1.6% (ICC = 0.016) and hour of day only 0.3% (ICC = 0.003). Estimated residual variance (unaccounted by any of these groupings) was 69.7%, attributable to camera settings, flash, angle, zoom level, and substrate variation within individual photos.

**Table 3.** ICC variance decomposition.

| Source | ICC | % Variance | N Groups | N Obs |
| --- | --- | --- | --- | --- |
| Observer (user_id) | 0.233 | 23.3% | 33,496 | 103,653 |
| Geography (H3 res-5) | 0.051 | 5.1% | 6,474 | 103,653 |
| Species | 0.016 | 1.6% | 34 | 103,653 |
| Hour of day | 0.003 | 0.3% | 24 | 103,653 |
| Residual | — | 69.7% | — | — |

*Note: ICCs were estimated from separate one-way ANOVAs; grouping variables are not orthogonal (e.g., observer and geography are correlated). The residual is defined as 100% minus the sum of individual*
ICCs. A single multilevel model with crossed random effects would yield a different decomposition. The mixed model estimate of 25.8% for observer (below) provides a partial correction.

The linear mixed model with `user_id` as a random intercept confirmed the observer variance estimate: the photographer random effect accounted for 25.8% of total brightness variance, whereas the fixed effects collectively added little explanatory power beyond the grand mean. Hour-of-day analysis showed a modest brightening trend from dawn through midday followed by decline through evening, but the ICC of 0.003 indicates that time-of-day effects are minor relative to observer identity. The distribution of per-observer mean brightness (among observers with ≥5 observations) had a standard deviation of ∼16.7 V units on a 0–255 scale, confirming that photographers introduce systematic additive offsets amounting to roughly 6.5% of the full brightness range.

### Crop Quality Validation and Sensitivity Analysis

Manual scoring of 200 audited crops revealed that only 76 (38.0%) were clearly acceptable for dorsal-body brightness extraction (`crop_quality = good`; Figure 5A-B; Table 4). The remaining 62.0% were `partial` (n = 58; 29.0%), `fail` (n = 24; 12.0%), or `in_hand` (n = 42; 21.0%). The automated QC filter (`passed_qc`: brightness □ [15, 245], entropy ≥ 4.0) failed to reject a single image in the 200-image audit sample: 100% of images across all four quality categories, including all 42 `in_hand` images, passed the automated screen (Figure 5C). This confirms that `passed_qc` functions as a data-quality filter (rejecting overexposed or uniformly dark images) rather than a localization check.

**Table 4.** Manual crop quality audit (n = 200 images).

| Label | n | % | brightness_usable | morph_usable |
| --- | --- | --- | --- | --- |
| `good` | 76 | 38.0% | yes | yes |
| `partial` | 58 | 29.0% | unclear | yes |
| `fail` | 24 | 12.0% | no | no |
| `in_hand` | 42 | 21.0% | no | unclear |

**Table 5.** Sensitivity analysis comparison.

| Dataset | n (obs) | Brightness $R^2$ | Morph $R^2$ (cells) |
| --- | --- | --- | --- |
| Full dataset | 103,653 | 0.0011 | 0.0080 |
| Path A: audit-clean subset | 75 | 0.027 ($p = 0.37$) | 0.360 ($p = 0.51$; 6 cells) |
| Path C: classifier-filtered | 10,823 | 0.0008 | 0.022 ($p = 0.06$) |

**Figure 5.**
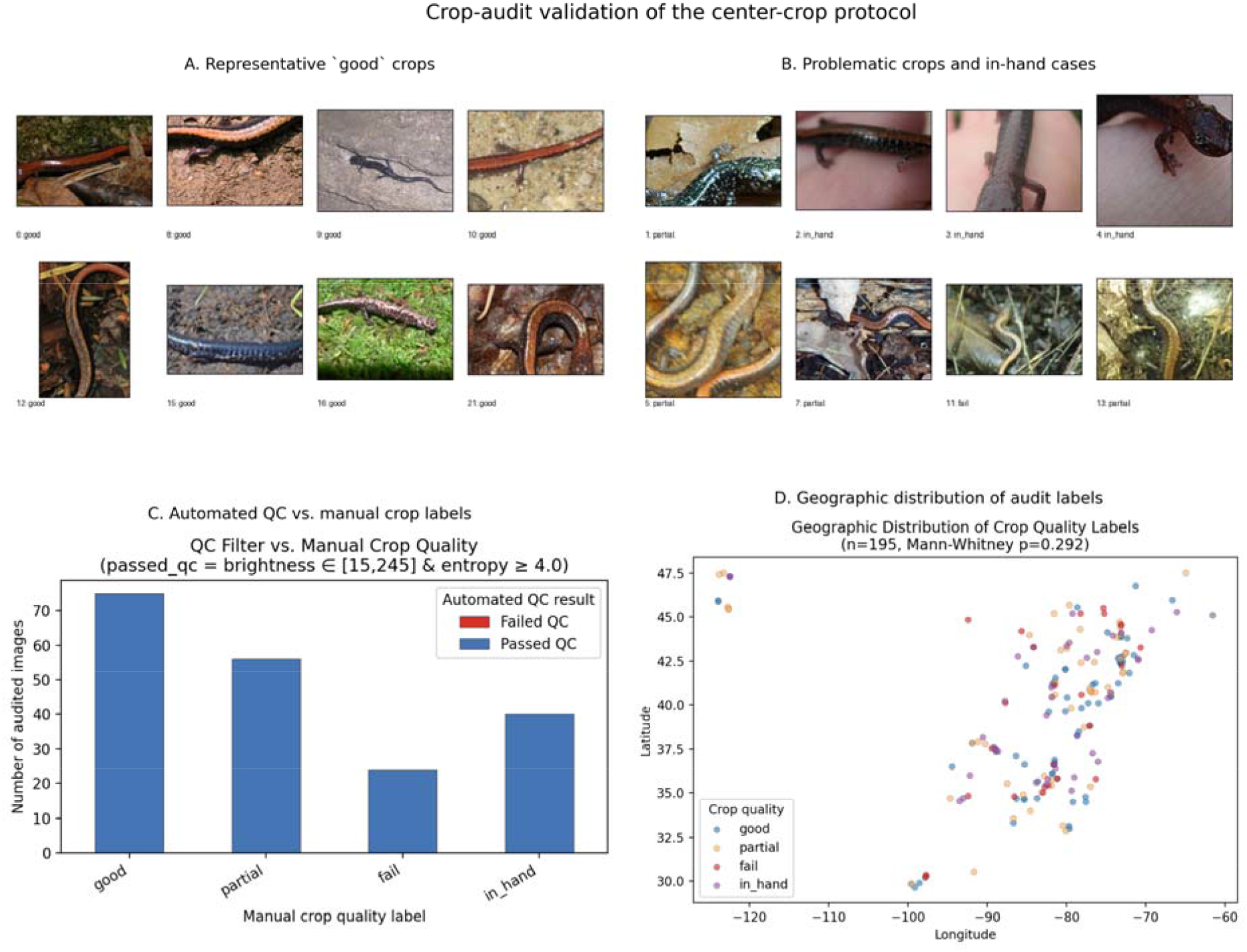
Crop-audit validation of the center-crop protocol.(A) Representative audited crops scored ‘good’. (B) Representative problematic crops, including partial-body and in-hand cases. (C) Concordance between automated ‘passed_qc’ and manual crop-quality labels. (D) Geographic distribution of audited crop-quality labels, showing no strong latitudinal bias in crop failures.

The latitude distributions of `good` and `not-good` crops were not significantly different (Mann–Whitney U = 4,096, p = 0.29; good mean lat = 39.3°, bad mean lat = 40.0°; Figure 5D), indicating that crop failures are geographically random rather than systematically concentrated at particular latitudes. This supports treating bad crops as additive measurement noise rather than a source of directional geographic bias.

#### Path A (audit clean subset)

Restricting to the 75 observations scored `brightness_usable = yes`, OLS regression of brightness ∼ lat + lon returned R^2^ = 0.027 (n = 75, p = 0.37). The estimate is not statistically significant and is consistent with the null result from the full dataset (R^2^ = 0.001), though the small sample precludes strong inference. On the 67 *P. cinereus* observations scored `morph_usable = yes`, the ambiguous classification rate dropped to 16.4% (vs. 20.3% in the full dataset), consistent with the interpretation that most ambiguous calls originate from in-hand photographs. A cell-level morph regression on the classified subset returned R^2^ = 0.360 across only 6 cells (p = 0.51), which is too sparse to support strong inference despite the larger point estimate.

#### Path C (classifier-filtered full dataset)

A logistic regression classifier trained on 195 audit-labeled observations achieved 5-fold cross-validation accuracy of 0.65 (majority-class baseline = 0.39), indicating modest above-chance discriminability. Applied to the full 103,653-observation dataset, the classifier retained 10,823 observations (10.4%). Brightness R^2^ on the filtered subset was 0.0008 (n = 10,823), essentially identical to the full-dataset result. The classifier introduced a notable morph-sampling bias: the predicted-good *P. cinereus* subset was 67.5% red-back versus 49.2% in the predicted-bad subset, reflecting the spectral overlap between human skin-tone hues and red-back dorsal coloration in HSV space. Morph frequency R^2^ on the classifier-filtered subset was 0.022 (n = 255 cells, p = 0.06) — numerically higher than the full-dataset result (0.008) but not statistically significant, and likely inflated by the morph-composition bias rather than reflecting genuine improvement in localization quality.

### Color Morph Classification and Geographic Signal

The hue-threshold classifier applied to 71,627 *P. cinereus* observations classified 29,004 (40.5%) as red-back, 28,062 (39.2%) as lead-back, and 14,561 (20.3%) as ambiguous. The 20.3% ambiguous rate is closely consistent with the 21.0% `in_hand` rate observed in the manual crop audit, indicating that most unclassifiable observations reflect photographic rather than phenotypic ambiguity: crops dominated by hand or glove pixels lack the hue/saturation signal needed to resolve morph identity. Red-back frequency across 3,746 H3 cells with ≥1 classified observation was 0.51 (mean across cells). Aggregating to cells with ≥5 classified observations (n = 1,776), OLS regression of red-back frequency ∼ mean cell latitude + mean cell longitude returned R^2^ = 0.008 (lat-only R^2^ = 0.003; F-test p < 0.001; Figure 1B, 1C). This is 7× larger than the brightness R^2^ from the same photographs (0.008 vs. 0.001), and statistically significant, confirming that a detectable geographic signal in morph composition exists in these photos — one that the continuous brightness analysis completely fails to recover.

The morph map (Figure 1B) shows spatially coherent variation in red-back frequency across the *P. cinereus* range, with a modest northward and westward increase in red-back frequency consistent with published regional and range-wide surveys (Gibbs and Karraker 2006; Hantak et al. 2019; Hantak et al. 2022; Moore and Ouellet 2015). However, classified red-back frequency (51%) falls substantially below published field-survey estimates for this species, which range from ∼74–80% range-wide (Gibbs and Karraker 2006) to >95% in northern populations (McCormick and Riley 2025), indicating that the classifier systematically underestimates red-back prevalence. I attribute this to observer novelty bias: lead-back individuals are photographed disproportionately relative to their population frequency because they appear unusual to citizen scientists, inflating the lead-back fraction of the iNaturalist sample (McCormick and Riley 2025).

## Discussion

### The Autoresearch Loop as a Phenotyping Optimization Tool

The *autoresearch* framework (Karpathy 2026) was designed for machine-learning hyperparameter search, in which a clear, automatable loss function evaluates candidate configurations within a constrained search space. Here, I show that the same architecture transfers naturally to ecological image-pipeline optimization, where the objective function combines geographic signal strength with within-cell measurement noise. The 50-experiment pilot loop identified four actionable improvements, namely CIE L\*a\*b\* color space, histogram normalization, a larger crop fraction, and moderate percentile trimming, that reduced within-cell variance by 97% while leaving the latitude R^2^ unchanged. This outcome illustrates a useful property of the approach: a well-specified metric can distinguish changes that improve precision from changes that merely amplify noise or compress signal.

The 24% acceptance rate is also informative about the optimization landscape. Most parameter perturbations either failed to improve the composite score or traded signal for noise reduction. Background masking, although initially appealing as a way to exclude leaf litter and substrate pixels from the crop, never improved the metric, likely because the centered crop already excluded most background and masking introduced edge artifacts that inflated within-cell variance. That conclusion would have been difficult to reach through informal manual iteration; the loop established it quickly and transparently. More broadly, the *autoresearch* pattern should be useful for phenotyping pipelines in which (1) the parameter space is sufficiently bounded for sequential search, (2) a composite evaluation metric encoding both sensitivity and reproducibility can be formalized, and (3) per-experiment cost is low enough to permit many iterations. The ∼$21 total cloud cost for the full production run, including a preemption-recovery restart, suggests that this approach is feasible without unusually large compute budgets.

### Why Continuous Brightness Fails: The Noise Floor

The primary geographic analysis returned a well-powered null result under the present measurement approach: brightness ∼ latitude + longitude explains R^2^ = 0.001 across 103,653 observations and 34 species, and the same non-result holds within *P. cinereus* alone across 71,627 individuals spanning the species’ full geographic range. This is not a statistical power problem — at n = 71,627, this provides >99% power to detect R^2^ ≥ 0.01. The result indicates that no detectable geographic brightness cline emerges from these photos once measurement noise is taken into account.

The variance decomposition explains this result. Observer identity accounts for 23.3% of brightness variance, a value corroborated by the mixed model estimate of 25.8%. The 33,496 photographers represented in this dataset introduce characteristic brightness offsets on the order of ±16.7 V units among observers with at least five observations, plausibly reflecting device-specific exposure behavior, flash use, and preferred shooting distance. Any real latitudinal gradient in dorsal brightness would therefore need to exceed this observer noise floor to be recoverable from raw pixel values. At the 0–255 scale used here, the observed inter-observer offsets imply that only comparatively large geographic shifts in mean brightness would be detectable.

The remaining 69.7% residual variance captures additional uncontrolled heterogeneity in citizen-science photography, including viewing angle, zoom level, substrate leakage into the crop, and background illumination. The geographic cell ICC (5.1%) is larger than the species ICC (1.6%), suggesting that locations differ not only biologically but also in the characteristic conditions under which observations are recorded. Hour of day (0.3% variance) proved comparatively unimportant, likely because most *Plethodon* observations are made during evening and nighttime coverboard searches, which restrict the range of ambient lighting conditions.

A manual crop quality audit of 200 images provides direct evidence that a substantial fraction of this residual variance originates from localization failures. Only 38% of audited crops were clearly suitable for dorsal brightness extraction; 21% showed the animal in hand, and an additional 41% were partial or unusable frames. Critically, the automated QC filter (brightness + entropy thresholds) did not reject a single image in the audit sample — confirming that `passed_qc` is not a localization check. Two sensitivity analyses support the robustness of the null brightness result. Restricting to the 75 audit-verified clean observations returned a non-significant R^2^ = 0.027 (p = 0.37), consistent with the null. A logistic regression classifier applied to the full 103K dataset retained 10,823 predicted-good observations and returned R^2^ = 0.0008. A geographic bias test showed no significant difference in latitude between good and poor-quality crops (Mann–Whitney p = 0.29), indicating that crop failures are randomly distributed across the range. Random localization failures inflate measurement noise but cannot create a geographic brightness cline where none exists; the null result is therefore not an artifact of the crop failure rate.

The optimized extraction parameters — histogram normalization especially — reduced within-cell variance by 97% in the pilot, but this compression of photographic noise did not reveal a previously hidden geographic signal. This is the critical diagnostic: if photographic noise were masking a real signal, normalization would have uncovered it by improving the signal-to-noise ratio. The fact that R^2^ remained stable at ∼0.018 (pilot) and 0.001 (full dataset) despite dramatically reduced noise indicates that the geographic brightness signal in iNaturalist *Plethodon* photos is either absent or smaller than any photometric normalization can recover.

### What Discrete Morphs Can Detect

The hue-threshold morph classifier recovered a geographic signal in *P. cinereus* red-back frequency (R^2^ = 0.008) that is 7× larger than the brightness R^2^ from the same photos. This serves as a narrow positive control: the same photographs that fail to reveal a continuous brightness gradient still contain sufficient color information to classify morphs and detect geographic variation in morph frequency. The signal is modest (R^2^ = 0.008 at the cell level), but statistically significant and directionally consistent with published range-wide surveys (Gibbs and Karraker 2006; Moore and Ouellet 2015).

However, direct comparison with Hantak et al. (2022) reveals the limits of the unsupervised threshold approach. Their ensemble CNN, trained on 4,000 human-labeled images, achieved ∼98% classification accuracy and recovered climatic associations with pseudo-R^2^ ≈ 0.04 — roughly 5× stronger than the present threshold-based result from the same image source (though note that OLS R^2^ and pseudo-R^2^ are not directly comparable metrics, so this ratio is approximate). The performance gap is attributable to several factors. First, the CNN implicitly localizes the dorsal stripe by learning to attend to the relevant image region, whereas the central crop approach averages over the entire frame including irrelevant areas. Second, the CNN’s whole-image representation is robust to common failure modes (blur, partial occlusion, oblique angles) that degrade mean hue and saturation measurements. Third, the CNN was trained specifically on this classification task, encoding expert knowledge about what constitutes “red-back” versus “lead-back” appearance under real photographic conditions. The threshold classifier embeds a simplified model of morph appearance (warm hue + adequate saturation) that holds for well-lit, dorsum-visible photos but degrades under adverse conditions, producing the conservative 40.5%/39.2% red-back/lead-back split (versus 75.9%/24.1% in Hantak et al.) and an ambiguous fraction of 20.3%. The crop audit provides a direct explanation for this ambiguous rate: 21.0% of audited images were scored `in_hand`, and crops dominated by hand or glove pixels lack the color signal needed to resolve morph identity. The CNN approach of Hantak et al. (2022) does not produce an analogous ambiguous class because the whole-image deep learning architecture implicitly localizes the dorsal region and is trained to classify every input; the threshold approach propagates crop failures directly into ambiguous calls.

The progression across methods, from continuous brightness (R^2^ = 0.001) to a simple hue threshold (R^2^ = 0.008) to a supervised CNN (pseudo-R^2^ ≈ 0.04), quantifies the signal-to-noise problem from a methodological perspective: more sophisticated classifiers extract more signal from the same photographs, but require correspondingly greater investment in training data and model development.

### Observer Novelty Bias and Sampling Distortion

McCormick and Riley (2025) documented that iNaturalist observations of *P. cinereus* in New Brunswick overrepresent rarer morphs relative to concurrent field surveys (unstriped: 6.6% on iNaturalist vs. 3.3% in the field; χ^2^ = 5.83, p = 0.02), attributing this to citizen scientists’ preference for photographing visually unusual individuals. The present dataset shows a related pattern: even after conservative morph classification, 39.2% of classified observations are lead-back. Published estimates of unstriped morph frequency vary substantially across the species’ range — from <5% in many northern populations (Lotter and Scott 1977; McCormick and Riley 2025) to 20–26% range-wide (Gibbs and Karraker 2006, compiling 50,960 individuals from 558 sites; see also Hantak et al. 2021) — but the 39.2% observed here substantially exceeds even the highest published field estimates. Moore and Ouellet (2015), analyzing 236,109 observations from 1,148 localities, found no significant climatic or geographic influence on morph proportions, suggesting that the geographic pattern itself remains contested. Hantak et al.’s (2022) more accurate CNN classifier obtained 24.1% lead-back in their iNaturalist dataset — still above field survey estimates, confirming that the bias operates at the sampling (who gets photographed) rather than the classification (who gets correctly identified) level.

This sampling bias has direct consequences for geographic inference. If the probability that an observer photographs an unusual morph varies by region — for example, if birding communities in the northeastern United States are more attentive to unusual herps than communities in the mid-Atlantic — then morph frequency estimates from iNaturalist will reflect photographer demographics as well as true biology. The geographic cell ICC from the variance decomposition (5.1%) is consistent with this: part of what looks like “geographic” brightness variation is likely geographically structured photographer behavior. The morph frequency R^2^ = 0.008 should therefore be interpreted cautiously; the modest geographic gradient may partly reflect spatial variation in observer novelty-seeking behavior rather than true morph frequency clines.

### Implications for Citizen Science Phenotyping

These results support a simple, practical framework for evaluating citizen science photos as phenotypic data sources. Continuous quantitative traits (brightness, size measurements, meristic counts) are vulnerable to observer variance because the measurement is a direct function of the raw pixel values, which are dominated by photographic conditions. Without standardized photography protocols — consistent backgrounds, calibrated color cards, known focal distances — observer identity will typically explain more variance in continuous trait estimates than any biological variable of interest. Photometric normalization can reduce within-photographer variance substantially, but cannot recover geographic signal that is smaller than the inter-photographer offset distribution.

Discrete categorical traits are more robust because classification thresholds can absorb photographic variation. Red-back and lead-back morphs differ dramatically in dorsal hue — warm orange-red versus desaturated gray-brown — a contrast that is perceptually and photometrically large enough to survive substantial photographic noise. However, as McCormick and Riley (2025) and the present data both demonstrate, the sampling process itself introduces a second layer of bias — observer preferences — that classification accuracy alone cannot correct. The implication is that citizen-science photo data may be most valuable for documenting the *existence* and *distribution* of discrete phenotypic variants rather than their *frequencies* or *quantitative values*.

The autoresearch loop provides a general way to determine where a given trait falls along this spectrum. By formalizing pipeline evaluation as a composite metric and running a bounded search, researchers can assess whether any parameter configuration extracts geographic signal above the noise floor before committing to large-scale analysis. In that sense, the negative result reported here, namely that no tested parameter configuration recovered a brightness cline in *Plethodon*, is itself informative.

### Future Directions

Three extensions would substantially strengthen the present analysis. First, adding a background-subtraction step using a pretrained segmentation model (e.g., SAM; Kirillov et al. 2023) would allow brightness extraction from confirmed dorsal pixels only, reducing the substrate-leakage component of residual variance. Given the magnitude of observer effects, this would likely improve precision more than reveal new geographic signal, but it would sharpen the inference. Second, a stratified subsampling design that selects one photo per observer per H3 cell would decorrelate observer identity from geography, reducing the observer ICC at the cost of sample size. At the estimated 23.3% observer ICC and 33,496 unique observers, such a dataset would still exceed 10,000 observations. Third, extending the Hantak et al. (2022) CNN framework to the full *Plethodon* genus, through species-specific or transfer-learned models for additional polymorphic species, would test whether the geographic morph signal observed in *P. cinereus* generalizes across the clade.

The methodological contribution of this study, namely the formalization of citizen-science image-pipeline optimization as a bounded, logged, multi-experiment search, is independent of the biological result. I recommend this approach as a standard component of research programs that seek to extract quantitative phenotypic data from opportunistically collected photographs.

## Data and Code Availability

Code for data acquisition, image processing, optimization, and analysis is archived at https://doi.org/10.5281/zenodo.19050224. Derived experiment logs and manuscript figures referenced here were generated from that repository. The observation metadata analyzed in this study were retrieved from iNaturalist through the iNaturalist API; the platform-level occurrence dataset is cited in the References as a general source description for iNaturalist records.

## Acknowledgments

I thank iNaturalist and its community of contributors for making this dataset possible, Alex Pyron for his critical review, and Jessica Nadler for her improvements on content.

## References

Brodsky, I., Friend, A. J., and the h3-py contributors. 2018. h3-py: Python bindings for H3, a hierarchical hexagonal geospatial indexing system. Uber Technologies. https://github.com/uber/h3-py

Clusella-Trullas, S., van Wyk, J. H., and Spotila, J. R. 2007. Thermal melanism in ectotherms. Journal of Thermal Biology 32:235–245. 10.1016/j.jtherbio.2007.01.013

Delhey, K. 2019. A review of Gloger’s rule, an ecogeographical rule of colour: definitions, interpretations and evidence. Biological Reviews 94:1294–1316. 10.1111/brv.12503

Di Cecco, G. J., Barve, V., Belitz, M. W., Stucky, B. J., Guralnick, R. P., and Hurlbert, A. H. 2021. Observing the observers: how participants contribute data to iNaturalist and implications for biodiversity science. BioScience 71:1179–1188. 10.1093/biosci/biab093

Endler, J. A. 1980. Natural selection on color patterns in Poecilia reticulata. Evolution 34:76–91. 10.1111/j.1558-5646.1980.tb04790.x

Gibbs, J. P., and Karraker, N. E. 2006. Effects of warming conditions in eastern North American forests on red-backed salamander morphology. Conservation Biology 20:913–917. 10.1111/j.1523-1739.2006.00375.x

Hairston, N. G. 1951. Interspecies competition and its probable influence upon the vertical distribution of Appalachian salamanders of the genus Plethodon. Ecology 32:266–274. 10.2307/1930418

Hantak, M. M., Page, R. B., Converse, P. E., Anthony, C. D., Hickerson, C. M., and Kuchta, S. R. 2019. Do genetic structure and landscape heterogeneity impact color morph frequency in a polymorphic salamander? Ecography 42:1383–1394. 10.1111/ecog.04534

Hantak, M. M., Federico, N. A., Blackburn, D. C., and Guralnick, R. P. 2021. Rapid phenotypic change in a polymorphic salamander over 43 years. Scientific Reports 11:22681. 10.1038/s41598-021-02124-2

Hantak, M. M., Guralnick, R. P., Zare, A., and Stucky, B. J. 2022. Computer vision for assessing species color pattern variation from web-based community science images. iScience 25:104784. 10.1016/j.isci.2022.104784

Highton, R., Maha, G. C., and Maxson, L. R. 1989. Biochemical evolution in the slimy salamanders of the Plethodon glutinosus complex in the eastern United States. Illinois Biological Monographs 57:1– 153.

Highton, R. 1995. Speciation in eastern North American salamanders of the genus Plethodon. Annual Review of Ecology and Systematics 26:579–600. 10.1146/annurev.es.26.110195.003051

iNaturalist contributors, iNaturalist. 2026. iNaturalist Research-grade Observations. Occurrence dataset. Accessed 2026-03-12. 10.15468/ab3s5x

Jaeger, R. G. 1971. Competitive exclusion as a factor influencing the distributions of two species of terrestrial salamanders. Ecology 52:632–637. 10.2307/1934151

Karpathy, A. 2026. autoresearch. GitHub repository. Accessed 2026-03-12. https://github.com/karpathy/autoresearch

Kirillov, A., Mintun, E., Ravi, N., Mao, H., Rolland, C., Gustafson, L., Xiao, T., Whitehead, S., Berg, A. C., Lo, W.-Y., Dollar, P., and Girshick, R. 2023. Segment anything. 2023 IEEE/CVF International Conference on Computer Vision (ICCV), 3992–4003. 10.1109/ICCV51070.2023.00371

Lotter, F., and Scott, N. J., Jr. 1977. Correlation between climate and distribution of the color morphs of the salamander Plethodon cinereus. Copeia 1977:681–690. 10.2307/1443166

McCormick, A., and Riley, J. L. 2025. Integrating ecological and community science data to understand patterns of colour polymorphism and social behaviour at the northern range limit of a plethodontid salamander. PLOS ONE 20(9):e0332501. 10.1371/journal.pone.0332501

McLean, C. A., and Stuart-Fox, D. 2014. Geographic variation in animal colour polymorphisms and its role in speciation. Biological Reviews 89:860–873. 10.1111/brv.12083

Moore, J.-D., and Ouellet, M. 2015. Questioning the use of an amphibian colour morph as an indicator of climate change. Global Change Biology 21:566–571. 10.1111/gcb.12744

Schmidgall, S., Su, Y., Wang, Z., Sun, X., Wu, J., Yu, X., Liu, J., Moor, M., Liu, Z., and Barsoum, E. 2025. Agent Laboratory: Using LLM agents as research assistants. Findings of the Association for Computational Linguistics: EMNLP 2025, 5977–6043. 10.18653/v1/2025.findingsemnlp.320

